# Transcriptomes across fertilization and seed development in the water lily *Nymphaea thermarum* (Nymphaeales) reveal dynamic expression of DNA and histone methylation modifiers

**DOI:** 10.1101/2021.04.04.438399

**Authors:** Rebecca A. Povilus, William E. Friedman

## Abstract

Studies of gene expression during seed development have been performed for a growing collection of species from a phylogenetically broad sampling of flowering plants (angiosperms). However, attention has mostly been focused on crop species or a small number of ‘model’ systems. Information on gene expression during seed development is minimal for those angiosperm lineages whose origins predate the divergence of monocots and eudicots. In order to provide a new perspective on the early evolution of seed development in flowering plants, we sequenced transcriptomes of whole ovules and seeds from three key stages of reproductive development in the waterlily *Nymphaea thermarum*, an experimentally-tractable member of the Nymphaeales. We first explore general patterns of gene expression, beginning with mature ovules and continuing through fertilization into early- and mid-seed development. We then examine the expression of genes associated with DNA and histone methylation – processes known to be essential for development in distantly-related and structurally-divergent monocots and eudicots. Around 60% of transcripts putatively homologous to DNA and histone methylation modifiers are differentially expressed during seed development in *N. thermarum*, suggesting that the importance of dynamic epigenetic patterning during seed development dates to the earliest phases of angiosperm evolution. However, genes involved in establishing, maintaining, and removing methylation marks associated with genetic imprinting show a mix of conserved and unique expression patterns between *N. thermarum* and other flowering plants. Our data suggests that the regulation of imprinting has likely changed throughout angiosperm evolution, and furthermore identifies genes that merit further characterization in any angiosperm system.

## Introduction

Fertilization and seed development are critical parts of the plant life cycle that involve extensive transcriptional reprogramming. Seed development in flowering plants (angiosperms) is of particular interest, as it uniquely involves two separate fertilization events that produce two distinct offspring. Double fertilization in angiosperms occurs when a pollen tube reaches a mature ovule and delivers two sperm cells into the female gametophyte. Each sperm cell fuses with one of the two female gametes, the egg cell and the central cell, to produce (respectively) the embryo and the embryo-nourishing endosperm. While endosperm does not necessarily persist past seed germination, it surrounds the embryo throughout seed development and is a crucial mediator of the relationship between an embryo and its maternal sporophyte.

Given that seeds, and specifically endosperm, are the cornerstone of human diets, there has been much effort to understand the dynamic transcriptional landscape of seed development in a variety of economically important plants (maize (Li 2014)(Chen 2014), rice (Gao 2013)(Xu 2012), soybean (Jones 2013), peanut (Zhang 2012), camelina (Nguyen 2013), *Brassica* (Gao 2014, Ziegler 2019)) or model systems (*Arabidopsis* (Girke 2000, Belmonte 2013)). Yet little information exists from within lineages whose origins predate the divergence of monocots and eudicots, hindering an understanding of the evolution of developmental processes that contribute to seed development. While this is related to the difficulty of working with the vast majority of species within these lineages (e.g., *Amborella*, Nymphaeales, Austrobaileyales, Chloranthales, Ceratophyllales, magnoliids, which are typically long-lived trees, shrubs, lianas, or aquatic plants), a genetically and experimentally tractable species from within one of the most-early diverging lineages has been identified (Povilus 2015). *Nymphaea thermarum* (Nymphaeales) is a minute waterlily with a relatively short generation time and a draft genome assembly and annotation (Povilus 2020) – as such, it is poised to help illuminate questions about the evolution of flowering plant reproduction.

A common thread has emerged from studies of seed development in a wide variety of angiosperms: epigenetic patterning and imprinting is important for seed development, and particularly for endosperm development (Haig 1991)(Haig 2013)(Gehring 2017)(Satyaki 2017). Imprinting is a phenomenon that results in alleles with identical nucleotide sequences that have different expression patterns, depending on which parent the allele was inherited from (a “parent-of-origin” effect). In flowering plants, imprinting is largely understood to occur via the establishment of DNA and histone methylation patterns during gamete and seed development. Because methylation of DNA or histones can affect how a locus is expressed, different epigenetic patterns established during the development of male and female gametes can mean that certain loci are expressed preferentially from the maternally-or paternally-inherited copy of the allele (Zilberman 2006). Imprinting has been noted to be particularly important for the ability of endosperm to function as a nutritional mediator between the embryo and maternal sporophyte (Haig 1991)(Gehring 2017). However, some of the mechanisms that control DNA or histone methylation patterns appear to differ between monocots and eudicots (Furihata 2016)(Nalaamilli 2013)(Köhler 2012), leading to the question of when DNA and histone methylation, and their role in imprinting, became important aspects of seed development in flowering plants.

DNA methylation during reproductive development is perhaps best understood in *Arabidopsis and rice*, and involves the coordination of DNA methytransferases and demethylases, some of which operate as part of a RNA-dependent DNA methylation mechanism (Satyaki 2017). Members of the DNA METHYLTRANSFERASE family (MET) establish and maintain CG methylation, while CHROMOMETHYLASE proteins (CMT) establish or maintain CHG or CHH methylation. METs and CMTs are known to be expressed both in developing female gametophytes of *Arabidopsis*, as well as in offspring tissues after fertilization (Jullien 2012)(Köhler 2012). DEMETER (DME) is a DNA glycosylase that removes methylation established by MET1 and is active during female gametophyte development. DME activity determines expression of some of the components of the POLYCOMB REPRESSIVE COMPLEX 2 (PRC2) (Hsieh 2011). The PRC2 complex participates in histone H3K27 methylation, and in doing so regulates the expression of several genes known to be important for seed development (Hsieh 2011). PRC2 is comprised of MEDEA (MEA), FERTILIZATION-INDEPENDENT SEED2 (FIS2), FERTILIZATION-INDEPENDENT ENDOSPERM (FIE), and MULTICOPY SUPRESSOR OF IRA1 (MSI1), and is active in both the central cell of the female gametophyte and in endosperm (Furihata 2016). Together, METs, CMTs, DME, and PRC2 regulate methylation patterns necessary for imprinting of parent-or-origin specific expression patterns. Methylation-modifying processes not necessarily associated with imprinting, such as RNA-directed DNA methylation (RdDM), are also active during reproductive development and have been tied to repression of transposon activity in the egg cell, embryo, or other tissues (Köhler 2012)(Gehring 2017)(Ingouff 2017)(Satyaki 2017).

In order to shed light on whether imprinting via DNA and histone methylation could be responsible for recently discovered parent-of-origin effects on endosperm and embryo development in *N. thermarum* (Povilus 2018), we examined the expression patterns of genes involved in DNA and histone methylation, and in particular those known to be important for imprinting. By obtaining libraries of gene expression during important stages of seed development in the water lily *Nymphaea thermarum*, we provide the first such dataset from within any early-diverging angiosperm lineage. The three stages sampled represent unique suites of developmental processes (Figure 1)(Povilus 2015). The first stage of 0 DAA (days after anthesis) consists of whole, unfertilized ovules. The second stage is whole seeds at 7 DAA, when the endosperm is expanding but the embryo is relatively quiescent. Nutrients are actively being acquired by, and stored in, a tissue called the perisperm (a maternal sporophyte tissue derived from the nucellus). The third stage sampled is whole seeds at 15 DAA, when endosperm expansion and differentiation has nearly been completed. The embryo begins to undergo significant growth and morphogenesis, displacing space occupied by degenerating endosperm cells. The perisperm continues to acquire and store nutrient reserves. Thus, while seed components were not spatially dissected from each other, the selected time points capture important developmental landmarks for the embryo and endosperm.

**Figure 1:**
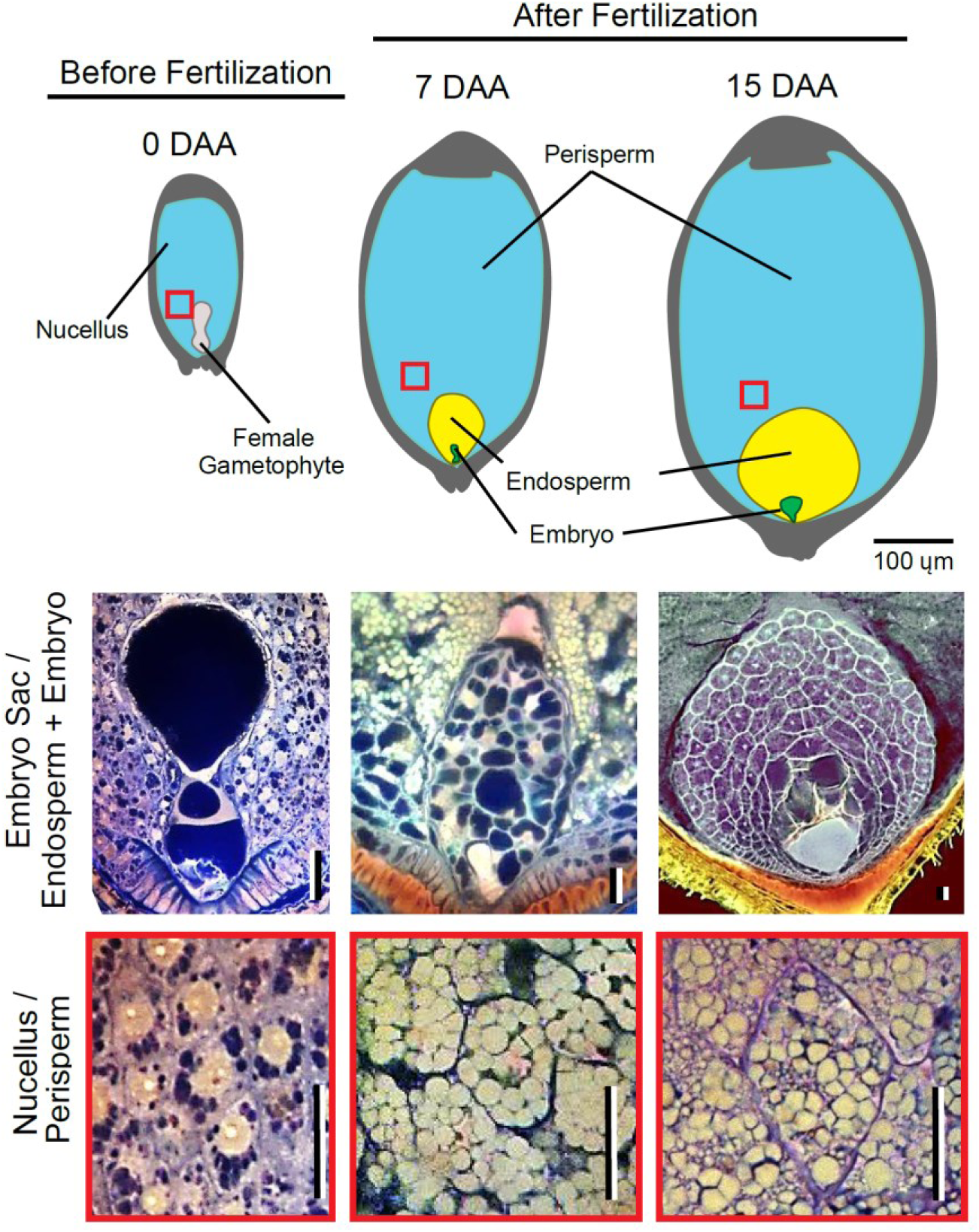
Stages of ovule and seed development in *N. thermarum* sampled for RNA-seq. Top row: Diagrams of the internal structure of ovules (before fertilization, 0 days before anthesis (DAA)) and seeds (after fertilization) at 7 DAA and 15 DAA, with key components labeled. Red boxes indicate location of corresponding image of the nucellus/perisperm (featured in the bottom row). Scale bar = 100 ųm. Middle and bottom rows: Confocal images of key ovule or seed components at each stage. Scale bars = 20 ųm.

We compare the expression profiles of genes that regulate DNA and histone methylation in *N. thermarum* with those of their homologs in monocots and *Arabidopsis*. In doing so, we identify processes that have likely been involved in seed development since the earliest stages of angiosperm evolution. We also find that that all of the molecular processes known to be involved in imprinting are indeed expressed before and/or after fertilization – suggesting that imprinting could occur in *N. thermarum*.

## Materials and methods

### Plant Material and Sequencing

The *Nymphaea thermarum* plants sampled for this study were grown in a greenhouse at the Arnold Arboretum of Harvard University (Boston, MA, United States) according to previously established protocols (Fischer 2010). Flowers were allowed to self-fertilize. All samples were collected between 10 and 11 am on their respective collection days. Ovules and seeds were quickly dissected from surrounding carpel or fruit tissue, weighed, then immediately frozen in liquid nitrogen, and stored at −80 C. For 0 DAA, each biological replicate represents material from 3-4 individual flowers from different plants. For 7 and 15 DAA, each biological replicate includes material from a single fruit.

RNA extractions were performed with a modified protocol, originally for use with maize kernels (Wang 2012)(Supplementary Materials and Methods 3). RNA seq libraries were prepared by the Whitehead Genome Sequencing Core, according to the manufacturer protocols of the Illumina Standard mRNA-seq library preparation kit (Illumina) using poly A selection, and were sequenced at the Baur Core of Harvard University to generate 125-bp, paired-end reads on a Illumina HiSeq Platform. All 12 libraries were multiplexed and sequenced on 3 lanes.

### Read mapping and differential expression analysis

For each sample, kallisto (Bray 2016) was used to pseudo-align reads to the *Nymphaea thermarum* genome (Povilus 2020) and quantify transcript abundances. 100 mapping bootstraps were performed, using default parameters for paired-end reads. Kallisto reports both estimated number of transcript reads per sample (EST), as well as transcript abundance per million reads (TPM, normalized for transcript length and number of reads per sample)(Supplemental Dataset 1). Transcripts that are differentially expressed (DE) between time points were identified using sleuth (Pimentel 2017)(Supplemental Dataset 2). The primary DE analysis modeled the effect of time, for all time points, on transcript abundance. Subsequent DE analysis was conducted for pair-wise comparisons between time points (for this analysis, multiple testing was accounted for by requiring transcripts to be significantly DE according to the primary DE analysis). Sleuth incorporates information from bootstraps performed by kallisto to estimate the inferential variance of each transcript; the adjusted variances were used to determine differential expression for each transcript. Transcripts were considered differentially expressed if time point (DAA) was a significant factor for transcript abundance, according to both a conservative likelihood ratio test and a Wald test (multiple-testing corrected p-value < 0.01).

### PCA and Clustering of Biological Replicates

Analyses were performed with R (version 3.4.0, (R Core Team 2017)). To assess similarity of biological replicates, EST counts for each transcript that was differentially expressed were centered and scaled (according to transcript means across all samples), using the scale function. Dimensional expression data was reduced to two dimensions by PCA using the prcomp function. K-means clustering within PCA space was performed by the kmeans function, with cluster number set to 3 (number chosen to reflect the number of sampled time points).

### Expression pattern cluster definition and analysis

Analyses were performed with R (version 3.4.0, (R Core Team 2017)). K-means clustering of gene expression patterns was performed with sample TPM values, using the kmeans function. The cluster number was set to 9, as the use of higher cluster numbers failed to identify additional unique expression patterns. Only transcripts that were differentially expressed were used for k-means clustering. The z-score was calculated for each gene per sample, using the scale function. Only genes whose expression patterns correlated with the average profile of each cluster (Pearson correlation > 0.9) were used in further analysis (Supplementary Dataset 3).

GO (molecular function) enrichment for each expression pattern cluster was performed with agriGO (Tian 2017), using *Arabidopsis thaliana* TAIR 10 annotation as the background, with hypergeometric or chi-squared tests (chi-squared was only performed if the query list had relatively few intersections with the reference list, and is noted separately), Yekutieli (FDR under dependency) multiple corrections testing adjustment, and significance level = 0.1. Putative *A. thaliana* homologs of *N. thermarum* transcripts were identified via BLASTX for each *N. thermarum* transcript against a database of all TAIR 10 *Arabidopsis* transcript amino acid sequences (downloaded from Phytozome (Goodstein 2012)), using the hit with the lowest e-value as the putative homolog match (e-value cutoff = 1e-15).

### Identification and analysis of transcription factors

Putative transcription factors (and their respective family type) were identified from the *Nymphaea thermarum* genome using the ‘Prediction’ tool available from the Plant Transcription Factor Database v4.0 (Jin 2017). Enrichment analysis for TF families among the set of DE TFs in each expression cluster was performed in R with Fisher’s exact test; adjusted p-values (FDR) <0.1 and <0.05 are noted.

### Identification of genes involved in histone and DNA methylation

To comprehensively identify putative homologs of genes known to be involved in regulation of DNA and chromatin methylation during seed development in other angiosperms, we estimated gene-family phylogenies for gene families of particular interest (CMT, MET, DME, components of the PCR2 complex). For each gene family, amino acid sequences for *Arabidopsis* members were aligned and used as the input for HMMER searches (e-value cutoff = 1e-15) (Eddy 2011) to identify putative homologs from genomes of *Physcomitrella patens (Physcomitrella patens* v3.3, DOE-JGI, http://phytozome.jgi.doe.gov/), *Amborella trichopoda* (Amborella Genome Project 2013), *Nymphaea thermarum* (Povilus 2020), *Aquilegia coerulea (Aquilegia coerulea* Genome Sequencing Project, http://phytozome.jgi.doe.gov/), *Oryza sativa* (Ouyang 2007), *Zea mays (*Hirsch 2016), *Arabidopsis thaliana* (Lamesch 2012), and *Solanum lycopersicum* (Tomato Genome Consortium 2012). The latest versions of all annotated genome datasets, except for *N. thermaurm*, were downloaded from Phytozome (Goodstein 2012). Putative homologs were also identified from within the de-novo assembled, immature-ovule and non-seed transcriptomes of *N. thermarum* (which includes tissues from roots, floral buds, leaves, and pre-meiotic ovules)(Povilus 2020). The amino acid sequences for the set of all putative homologs for a gene family were aligned with MUSCLE (Edgar 2004), alignments were manually trimmed to represent highly conserved regions, and phylogenetic tree estimation and bootstrapping (n=100) was performed with RAxML under the PROTGAMMAGTR amino-acid substitution model (Stamataki 2014). During further discussion we took a conservative approach, using a relatively broad definition as to which members were included in particular gene sub-families of interest.

All *Arabidopsis* genes annotated as being involved with either DNA methylation (GO:0006306) and histone methylation (GO:0016571) were collected using QuickGO (http://www.ebi.ac.uk/). Putative *N. thermarum* homologs of *A. thaliana* transcripts were identified via BLASTX for each *N. thermarum* transcript against a database of all TAIR 10 *Arabidopsis* transcript amino acid sequences (Lamesch 2012), using the hit with the lowest e-value as the putative homolog match (e-value cutoff = 1e-15).

### Histology and Microscopy

Material collected for microscopy was fixed in 4% v/v acrolein (Polysciences, New Orleans, Louisiana, USA) in 1X PIPES buffer (50 mmol/L PIPES, 1 mmol/L MgSO4, 5 mmol/L EGTA) pH 6.8, for 24 hours. Fixed material was then rinsed three times (one hour per rinse) with 1X PIPES buffer, dehydrated through a graded ethanol series, and stored in 70% ethanol. Samples were prepared for confocal microscopy and imaged according previously established protocols (Povilus 2015). Briefly: tissues were stained according to the Fuelgen method, and then infiltrated with and embedded in JB-4 glycol methacrylate (Electron Microscopy Sciences, Hatfield, PA, USA). Blocks were cut by hand with razor blades to remove superfluous tissue layers. Samples were mounted in a drop of Immersol 518f (Zeiss, Oberkochen, Germany) on custom well slides and imaged with a Zeiss LSM700 Confocal Microscope, equipped with an AxioCam HRc camera (Zeiss, Oberkochen, Germany). Excitation/emission detection settings: excitation at 405 and 488 nm, emission detection between 400-520 nm (Channel 1) and 520-700 (Channel 2).

## Results

### Generation and analysis of RNA-seq Data

Between 66 and 100 million high quality reads were generated for each sample, for a total of 940 million reads. 76.2% of the reads pseudo-mapped to the *Nymphaea thermarum* genome, and uniquely mapped reads were used to estimate normalized transcript abundance as TPM (transcripts per million)(Supplementary Table 1). Biological replicates of each time point clustered with each other (and not with samples of other time points) during PCA and k-means analysis of expression patterns of the 4000 most highly expressed transcripts, except for one sample of 0 DAA seeds that clustered instead with 7 DAA samples (Figure 2A). This sample was removed from further analysis. When PCA and k-means clustering were performed with the remaining 11 samples, samples clustered according to collection time point.

**Figure 2:**
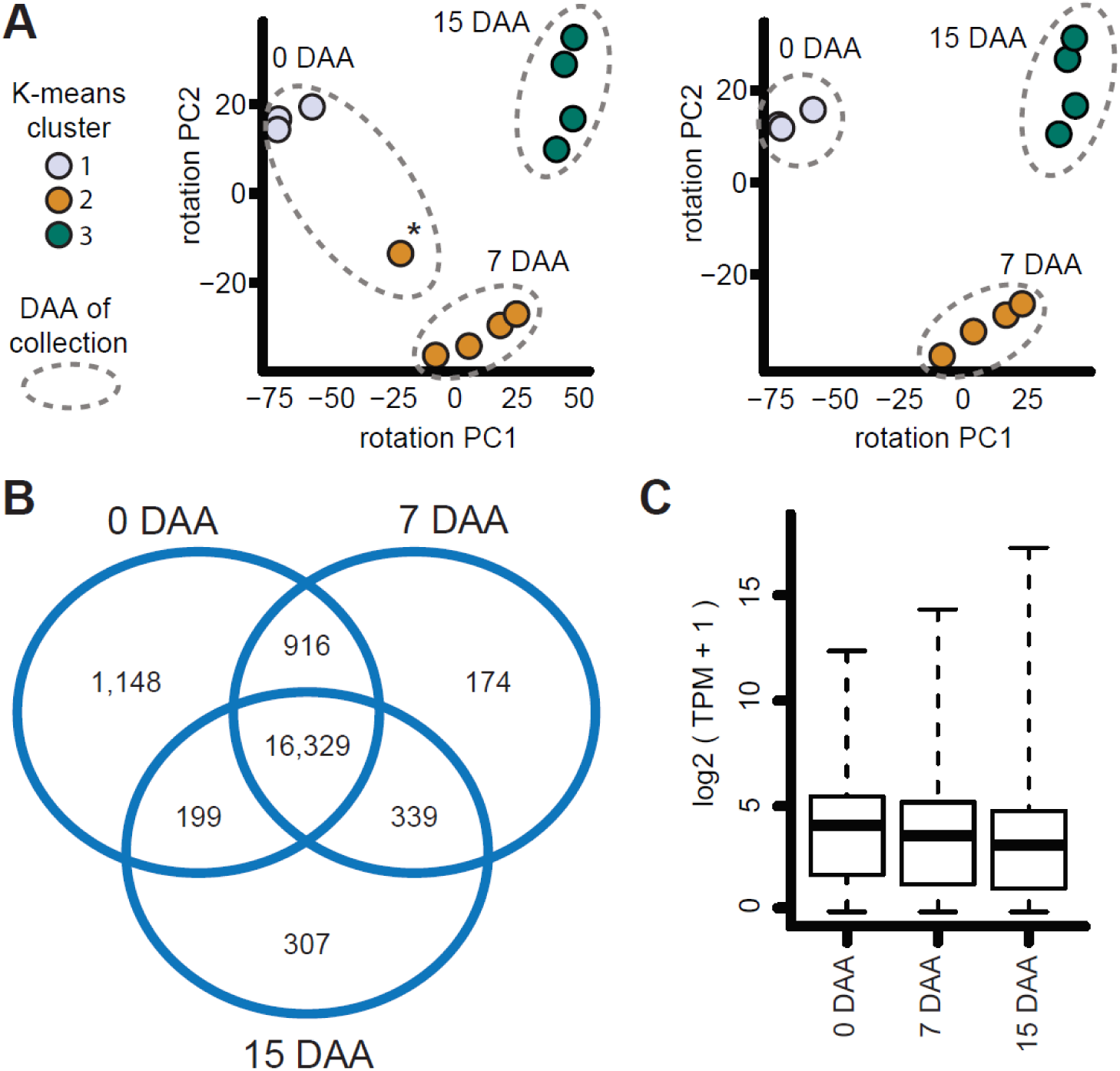
Basic Analysis of Transcriptomes. General characteristics of transcription in all samples. A) PCA of K-means clustering of biological replicates. Dot color indicates cluster identity, while inclusion within dashed outline indicates which DAA the sample was collected. Left graph: One 0 DAA sample clustered with 7 DAA samples (*). This sample was considered developmentally anomalous and removed from further analysis. Right graph: PCA without the anomalous sample. B) Venn diagram of unique transcripts with a TPM > 1 at each stage, and present in multiple stages. C) Distribution of TPM values for all transcripts (TPM >1) at each stage. Median values are indicated with bold horizontal lines, bottom and top of boxes indicate 25^th^ and 75^th^ percentile, and dashed lines indicate minimum and maximum values.

In total, 19,412 unique transcripts with at least a minimum abundance of 1 TPM were present during the sampled time points (Supplemental Dataset 1). This represents 74.4% of the 25,760 genes identified from the *Nymphaea thermaurm* genome (Figure 2B). 16,329 transcripts were present at all three ovule/seed developmental stages. 0 DAA had the most unique transcripts, and 7 DAA had the fewest. Among transcripts present in two stages, 0 DAA and 7 DAA shared the most transcripts, while 7 DAA and 15 DAA shared the fewest. The majority of transcript expression levels fell within a similar range across all three stages (Figure 2C). However, the expression levels of the 0.1% most highly expressed transcripts increased significantly between 7 and 15 DAA (Supplementary Table 2).

Besides differences in TPM values among the most highly expressed transcripts, the types of genes represented by the 10 most highly expressed transcripts varied with time point (Table 1). At 0 DAA, most of the 10 most highly expressed transcripts coded for structural components of histones or ribosomes. Notably, a putative homolog to an arabinogalactan peptide (AGP16) was the third most highly expressed transcript at 0 DAA. Arabinogalactans are known to regulate female gametophyte development and function in pollen tube interactions in other angiosperms (Pereira 2016). At 7 DAA, ribosome components again featured prominently among the 10 most highly expressed transcripts. A lipid transfer protein, WAXY starch synthase, and TPS10 terpene synthase were also highly expressed at 7 DAA, likely in relation to the initiation of nutrient import and storage in the seed, and seed coat differentiation. At 15 DAA, several transcripts involved with terpene synthesis or modification were among the 10 most highly expressed transcripts, coincident with continued maturation of the seed coat. A WAXY starch synthase, highly expressed at 7 DAA, continued to be highly expressed at 15 DAA.

**Table 1:** TPM, Identifier, and Puatative homology of the 10 transcripts with the highest abundances at each stage. Putative homology information was collected from the annotated genome of *N. thermarum* (Povilus 2020).

|  | TPM | Identifier | Putative Homology |
| --- | --- | --- | --- |
| 0DAA | 4966.98 | NYTH03818-RA | Ubiquitin (*Triticum aestivum*) |
|  | 4640.607 | NYTH03847-RA | Histone H4 (*Glycine max*) |
|  | 4466.02 | NYTH43515-RA | AGP16 Arabinogalactan peptide 16 (*Arabidopsis thaliana*) |
|  | 4220.917 | NYTH18212-RA | RPS30A 40S ribosomal protein S30 (*Arabidopsis thaliana*) |
|  | 3675.603 | NYTH26983-RA | Os01g0645000 Zinc finger CCCH domain-containing protein 9 (*Oryza sativa subsp. japonica*) |
|  | 3509.973 | NYTH40689-RA | RPL29A 60S ribosomal protein L29-1 (*Arabidopsis thaliana*) |
|  | 3436.017 | NYTH51439-RA | Defensin J1-2 (*Capsicum annuum*) |
|  | 3241.97 | NYTH00189-RA | At4g30220 Probable small nuclear ribonucleoprotein F (*Arabidopsis thaliana*) |
|  | 3104.7 | NYTH38977-RA | Protein of unknown function |
|  | 3056.18 | NYTH00233-RA | H2B Histone H2B.6 (*Arabidopsis thaliana*) |
| 7DAA | 19593.65 | NYTH51439-RA | Defensin J1-2 (*Capsicum annuum*) |
|  | 8724.83 | NYTH16912-RA | Non-specific lipid-transfer protein 1 (*Lens culinaris*) |
|  | 8283.093 | NYTH03818-RA | Ubiquitin (*Triticum aestivum*) |
|  | 6436.833 | NYTH18212-RA | RPS30A 40S ribosomal protein S30 (*Arabidopsis thaliana*) |
|  | 6126.765 | NYTH40689-RA | RPL29A 60S ribosomal protein L29-1 (*Arabidopsis thaliana*) |
|  | 6077.737 | NYTH45457-RA | HSP22 Small heat shock protein (*Glycine max*) |
|  | 4974.465 | NYTH18464-RA | WAXY Granule-bound starch synthase 1(*Antirrhinum majus*) |
|  | 4805.542 | NYTH27985-RA | Protein of unknown function |
|  | 4318.512 | NYTH59946-RA | TPS10 Terpene synthase 10 (*Ricinus communis*) |
|  | 4024.218 | NYTH38580-RA | RPL37C 60S ribosomal protein L37-3 (*Arabidopsis thaliana*) |
| 15DAA | 102890.9 | NYTH59946-RA | TPS10 Terpene synthase 10 (*Ricinus communis*) |
|  | 26734.5 | NYTH51439-RA | Defensin J1-2 (*Capsicum annuum*) |
|  | 18853.65 | NYTH44750-RA | (-)-alpha-terpineol synthase (*Vitis vinifera*) |
|  | 18529.12 | NYTH18464-RA | WAXY Granule-bound starch synthase 1(*Antirrhinum majus*) |
|  | 13610.54 | NYTH45457-RA | HSP22 Small heat shock protein, (*Glycine max*) |
|  | 9038.013 | NYTH16912-RA | Non-specific lipid-transfer protein 1 (*Lens culinaris*) |
|  | 8806.638 | NYTH28556-RA | Protein of unknown function |
|  | 8389.802 | NYTH59911-RA | Alpha-terpineol synthase, chloroplastic (*Magnolia grandiflora*) |
|  | 6565.69 | NYTH03818-RA | Ubiquitin (*Triticum aestivum*) |
|  | 6242.847 | NYTH59528-RA | TPS10 Terpene synthase 10 (*Ricinus communis*) |

### Analysis of Differential Gene Expression

Out of the 19,412 unique transcripts expressed during seed development in *N. thermarum*, 10,933 were significantly differentially expressed (DE) (Supplementary Dataset 2). The set of DE transcripts was used to perform hierarchical clustering of all transcripts in all samples (Figure 3A). The three main ‘clades’ identified by hierarchical clustering of samples correlated with the three sampling time points. 7 DAA samples and 15 DAA samples were more closely related to each other, than to 0 DAA samples.

**Figure 3:**
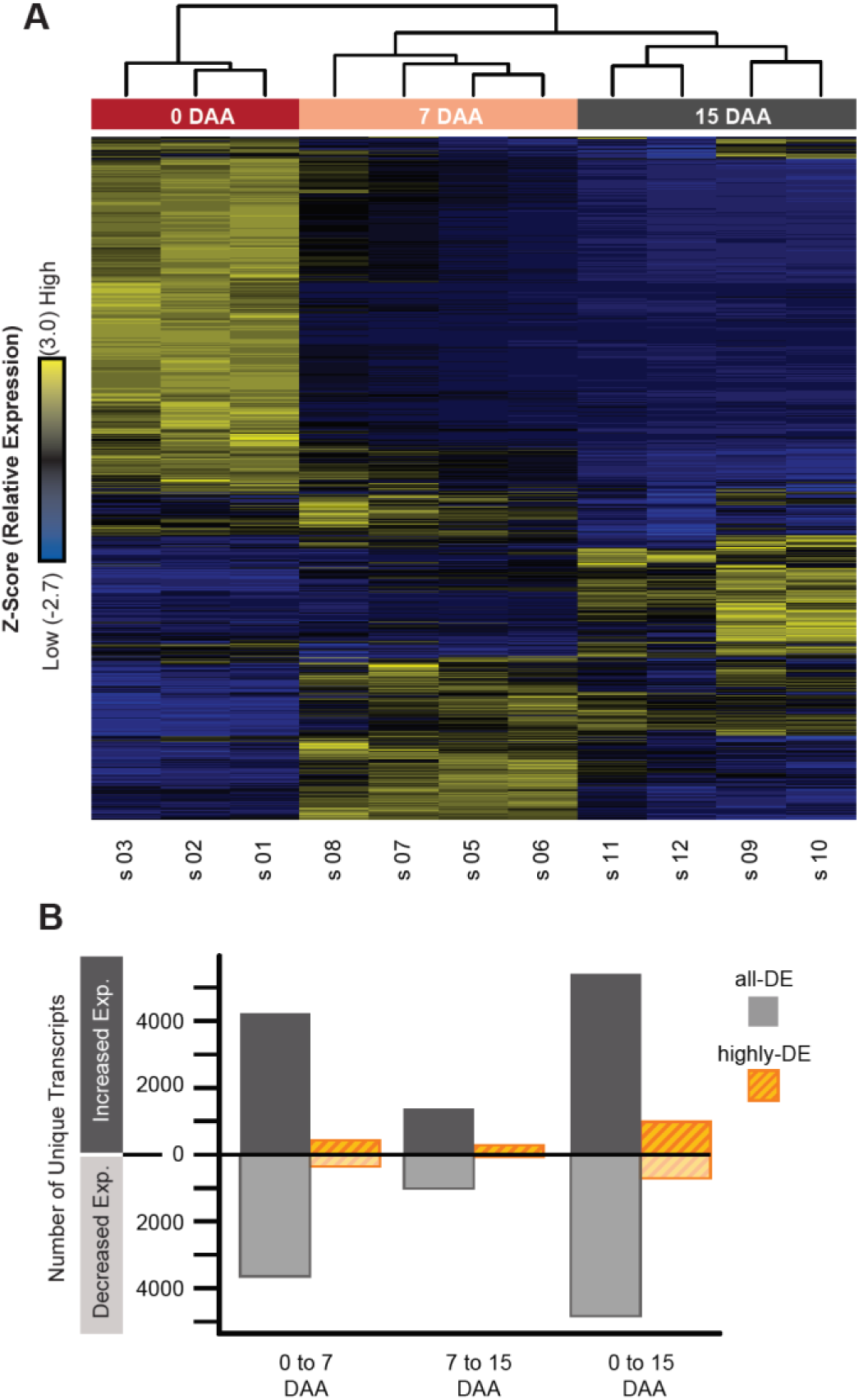
Differential Expression of Transcripts During Seed Development in *N. thermarum.* Basic analysis of differentially expressed (DE) transcripts. A) Heatmap of the relative expression (Z-scores of the mean TPM of all biological replicates at each stage) for each of 10,933 DE transcripts. Each row represents a single unique transcript; transcript identifies are not included. Rows are hierarchically clustered (dendrogram not included). Each column represents a single sample, and are hierarchically clustered (top dendrogram). B) Number of unique, DE transcripts that showed either an increase or decrease in expression between time points. Results from the set of all-DE or highly-DE transcripts are shown separately.

Two sets of differentially expressed (DE) transcripts were considered during further analysis: a set of all DE transcripts (“all-DE”), and a set of DE transcripts with very high relative changes in expression (“highly-DE”). For the latter set, an absolute b-value more than 2 for the expression change(s) was used as the filtering criteria. B-values are reported by sleuth (companion to the pseudo-mapping program kallisto) as part of differential gene expression analysis, and are analogous to fold-change in what a positive or negative value means for the direction of expression change (Pimentel 2017). However, b-values are derived from the effect size of time point on the log10-transformed transcript abundances – a b-value is therefore not equivalent to the same value fold-change (ie: a b-value of 2 does not imply a fold-change of 2).

Among the set of all DE transcripts (10,933 transcripts), more than three times more transcripts were differentially expressed between 0-7DAA, than between 7-15 DAA, while the number of transcripts differentially expressed between 0-7 DAA and 0-15 DAA was more similar (Figure 3B). For each time-point comparison, between 53 and 58% of the significant changes in expression were due to increases in expression (as opposed to decreases in expression). The set of highly-DE transcripts was smaller, consisting of 1,865 unique transcripts. The 7-15 DAA transition had the fewest highly-DE transcripts, with 0-7 DAA having about 2.5 times as many, and 0-15 DAA having about twice as many as 0-7DAA. While the proportion of transcripts that increased expression between 0-7DAA and 0-15 DAA was similar to what was seen in the set of all DE transcripts (between 54-58%), for 7-15DAA the proportion of highly-DE transcripts that increased expression was much higher (80%, as compared to 58% for the set of all DE transcripts).

### Analysis of transcripts grouped by expression pattern

The expression patterns of all DE transcripts were associated with 9 expression pattern types (or ‘clusters’) using a K-means clustering approach (Figure 4A) (Supplementary Table 3). The expression patterns represented by the 9 clusters include: increased or decreased expression across the entire time sampled (respectively, Clusters A and B), as well as increased or decreased expression to produce a minimum or maximum at each of the 3 time points (increase and decrease associated with minimum or maximum at, respectively, 0 DAA = Clusters C,D; 7 DAA = E,F; 15DAA = G,H). The final cluster (cluster I) represented transcripts that, while differentially expressed, displayed a relatively small magnitude of change. 10,450 transcripts (96% of DE transcripts) were strongly correlated with the average profile of their respective clusters (Pearson correlation > 0.9) and were considered for further analysis.

**Figure 4:**
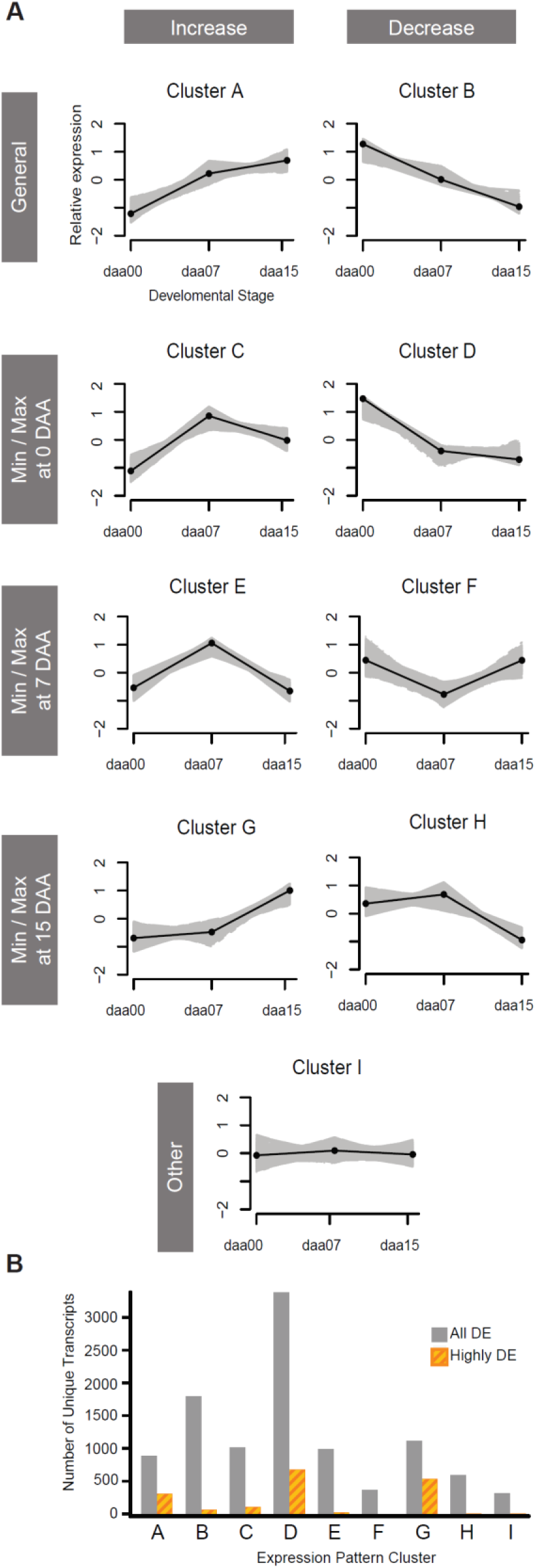
Expression Pattern Clusters for DE Transcripts. A) 9 expression pattern clusters, which contain 96% of all transcripts differentially expressed during the sampled stages of reproductive development. Mean TPM values of biological replicates at each stage were centered and scaled, relative to the mean transcript expression value over all stages. Clusters are organized by whether they represent initial increases or decreases (columns) to achieve consistent increased or decreased expression, or minimum or maximum expression at each stage (rows). Grey areas represent expression of each transcript in a cluster (only includes transcripts whose expression patterns correlated with the average profile of each cluster (Pearson correlation > 0.9)), while black lines represent the median expression pattern for each cluster. B) Number of unique transcripts in each cluster. Results from sets of all-DE and highly-DE transcripts are shown separately.

In addition to the set of 10,450 DE transcripts represented in the expression pattern clusters (“all-DE”), a subset of “highly-DE” transcripts (b-value > 2 for at least one time point transition: 1,783 transcripts) was considered during further analysis of expression clusters (Figure 4B). For the cluster pairs that represent general change (A,B) or minimum/maximum expression at 0 DAA (C,D), more transcripts showed expression decreases. In contrast, in the cluster pairs that represent minimum/maximum expression at 7 DAA (E,F) and 15 DAA (G,H), more transcripts showed increased expression.

### Functional enrichment of expression pattern clusters

Each cluster was tested for significant enrichment of GO molecular function terms, based on the TAIR 10 annotations for the putative *A. thaliana* homolog of each *N. thermarum* transcript. All significantly enriched child terms (ie: the most specialized of a hierarchy) are reported for each cluster, and terms of particular interest are further discussed (Figure 5).

**Figure 5:**
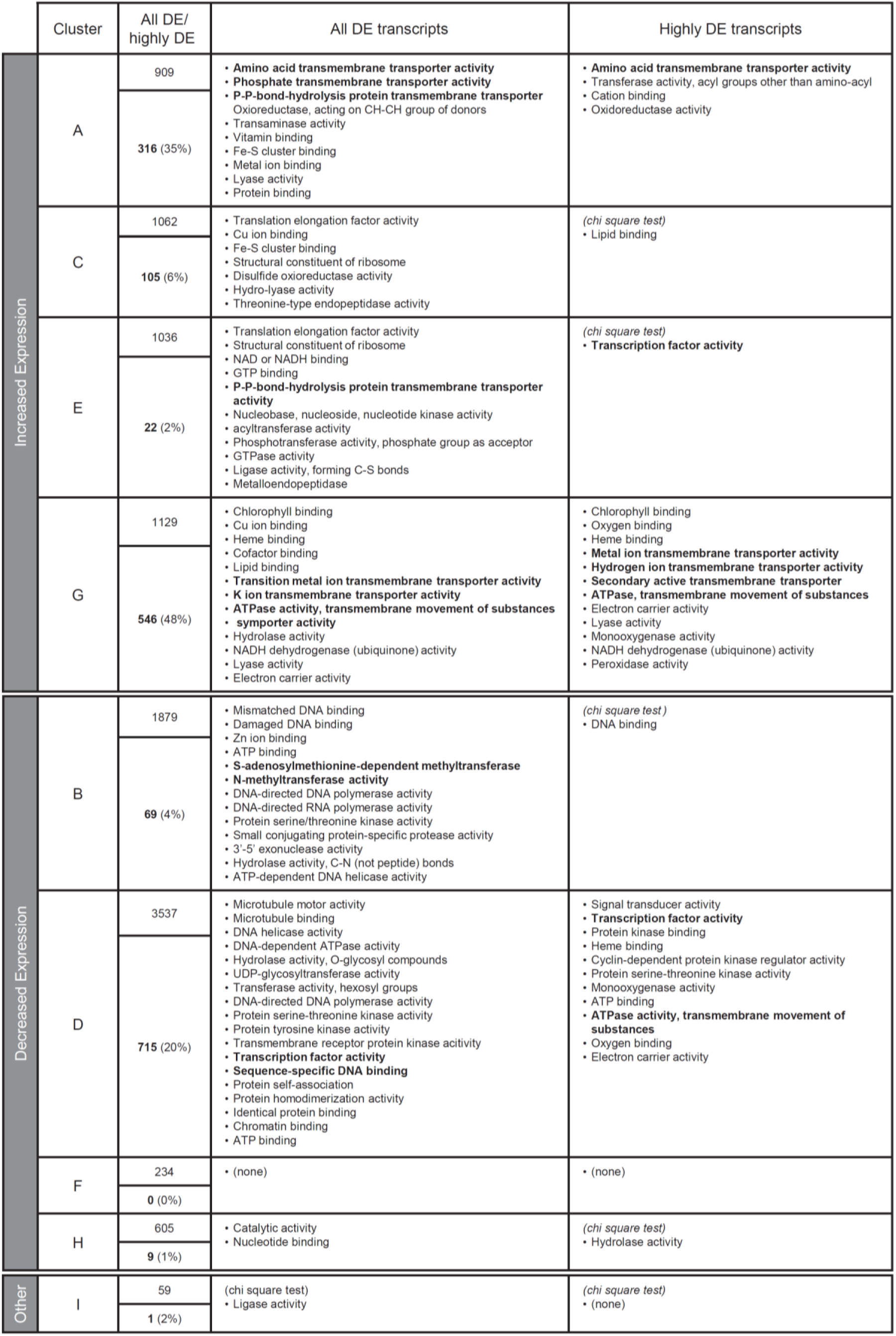
Summary of putative molecular functioned enriched in each expression pattern cluster. All significantly enriched child terms (ie: the most specialized of a hierarchy) are reported for each cluster. Unless otherwise noted, molecular function enriched was tested with hypergeometric test, using Yekutieli (FDR under dependency) multiple corrections testing adjustment, and significance level = 0.1. Molecular function in bold indicate functions of particular interest during discussion.

Clusters with general or time-point-specific increases in expression were found to be generally enriched for functions related to various types of transmembrane transporter activity. For the set of all-DE transcripts, Cluster G (maximum at 15 DAA) was additionally enriched for functions that appear to be related to chloroplast activity (chlorophyll-binding, electron-carrier activity). While it seems unlikely that seeds at this stage (which are enclosed within opaque fruit walls) would be carrying out photosynthesis, the analogous stage of embryo development in *Arabidopsis* is associated with the formation of chloroplasts within embryo tissues (Mansfield 1991). Among the set of highly-DE transcripts, various transmembrane transporter activities were again enriched in Clusters A (general increase) and G (maximum at 15DAA). Cluster C (increase after 0 DAA) was enriched for lipid binding, and Cluster E (maximum at 7DAA) was enriched for transcription factor activity.

Clusters associated with general or time-point-specific decreases in expression prominently featured significantly enriched terms related to DNA or chromatin binding and modification. For the set of all-DE transcripts, Cluster B (general decrease) was enriched for terms related to methyltransferase activity and Cluster D (decrease after 0 DAA) was enriched for transcription factor activity. Among the set of highly-DE transcripts, Cluster D (decrease after 0 DAA) was enriched for transcription factor activity.

### TF expression during seed development

The fact that clusters that represent both increases and decreases in expression were enriched for transcription factors merited further investigation of transcription factor activity. Of the 1,268 putative transcription factors identified from the *N. thermaurm* genome, 1,039 were expressed during the sampled stages (with a TPM > 1), and 719 were significantly differentially expressed. Among the set of DE transcription factors, we examined whether expression pattern clusters were significantly enriched for any of 58 transcription factor families (Figure 6). For the all-DE transcription factor dataset, Cluster B (general decrease) was enriched for FAR1, Cluster D (decrease after 0 DAA) was enriched for GRF, and Cluster E (maximum at 7 DAA) was enriched for MYB transcription factors. When only highly-DE transcription factors were considered (b > 2), Cluster C (increase after 0 DAA) was enriched for WRKY, Cluster D (decrease after 0 DAA) was enriched for ZF-HD and GRF, and cluster E (maximum at 7 DAA) was enriched for MYB activity.

**Figure 6:**
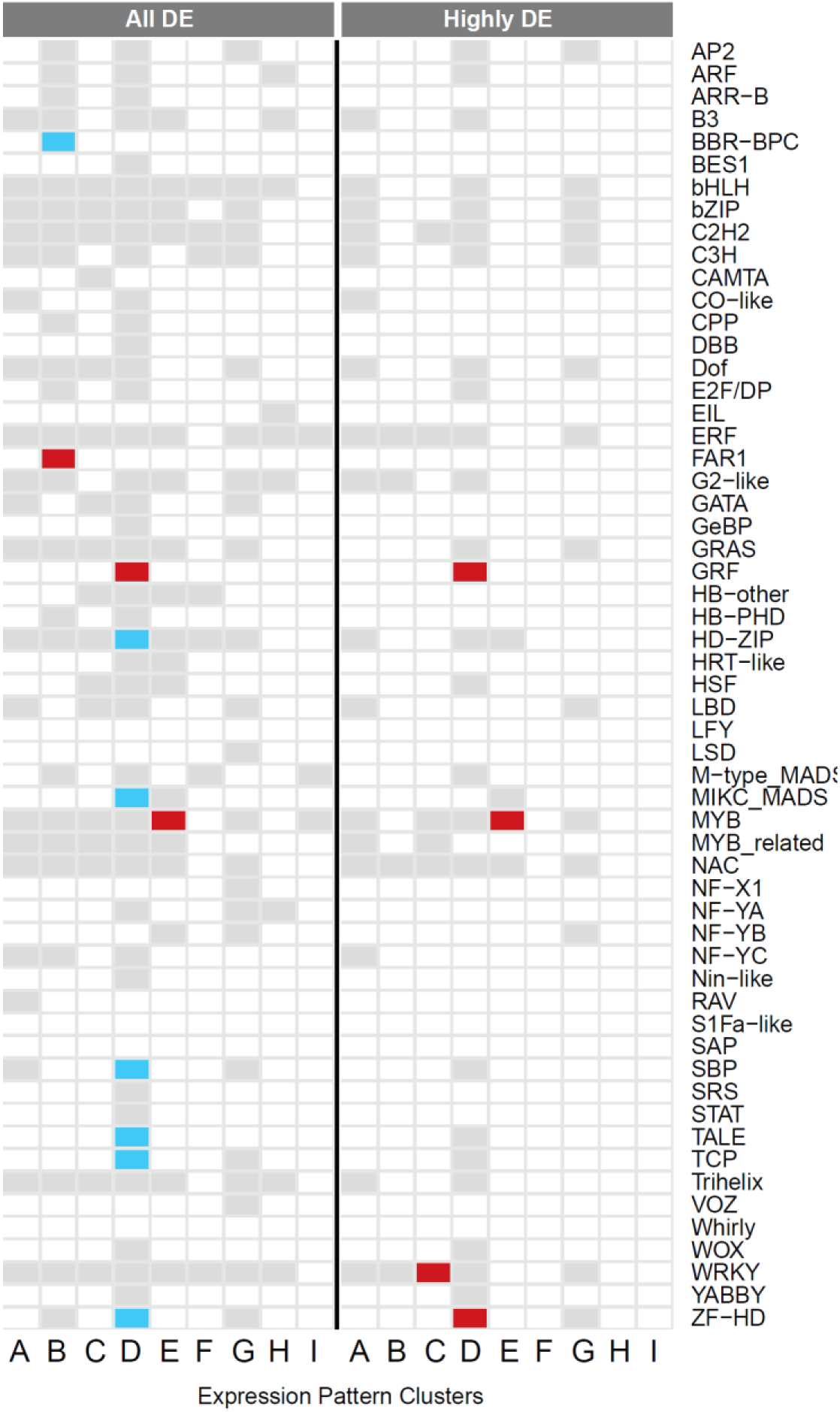
Enrichment of differentially-expressed transcription factor families. Enrichment analysis for TF families among the set of DE TFs in each expression cluster, performed with Fisher’s exact test; adjusted p-values (FDR) <0.1 (light blue) and <0.05 (dark red) are noted. Boxes in grey indicate at least one member of a TF family is present in an expression cluster, white indicates that no member of a TF family is present. Results for TFs from the sets of all-DE and highly-DE transcripts are reported separately.

### Activity of genes associated with imprinting via DNA and histone methylation

Methyltransferase-related terms were enriched in cluster B (consistent decrease in expression), already hinting at potential for a dynamic DNA and histone methylation landscape during reproductive development in *N. thermarum*. We constructed gene family phylogenies for genes that are known to be important regulators of epigenetic patterning during reproduction, with a particular focus on those involved in gene imprinting (CMT, MET, DME, and the PCR2 components MEA, FIS, FIE, and MSI). Many of the relationships between gene family members corroborates previous studies (Furihata 2016) (Bewick 2017).

First, we examined genes involved in the establishment or maintenance of imprinting-related DNA methylation in the CG cand CHG contexts: CMT and MET. Although there are 4 MET homologs in *Arabidopsis*, they appear to be the result of clade-specific gene duplications; the two *N. thermarum* MET homologs are similarly the result of a clade-specific gene duplication (Figure 7A). Only one *N. thermarum* CMT homolog was identified, although it’s affinity for either of the CMT2 or CMT1/CMT3 clades was poorly resolved (Figure 7B). All *N. thermarum* homologs of CMT and MET were differentially expressed during reproductive development in *N. thermarum*, and belonged to expression Cluster D (decrease after 0DAA) (Figure 9).

**Figure 7:**
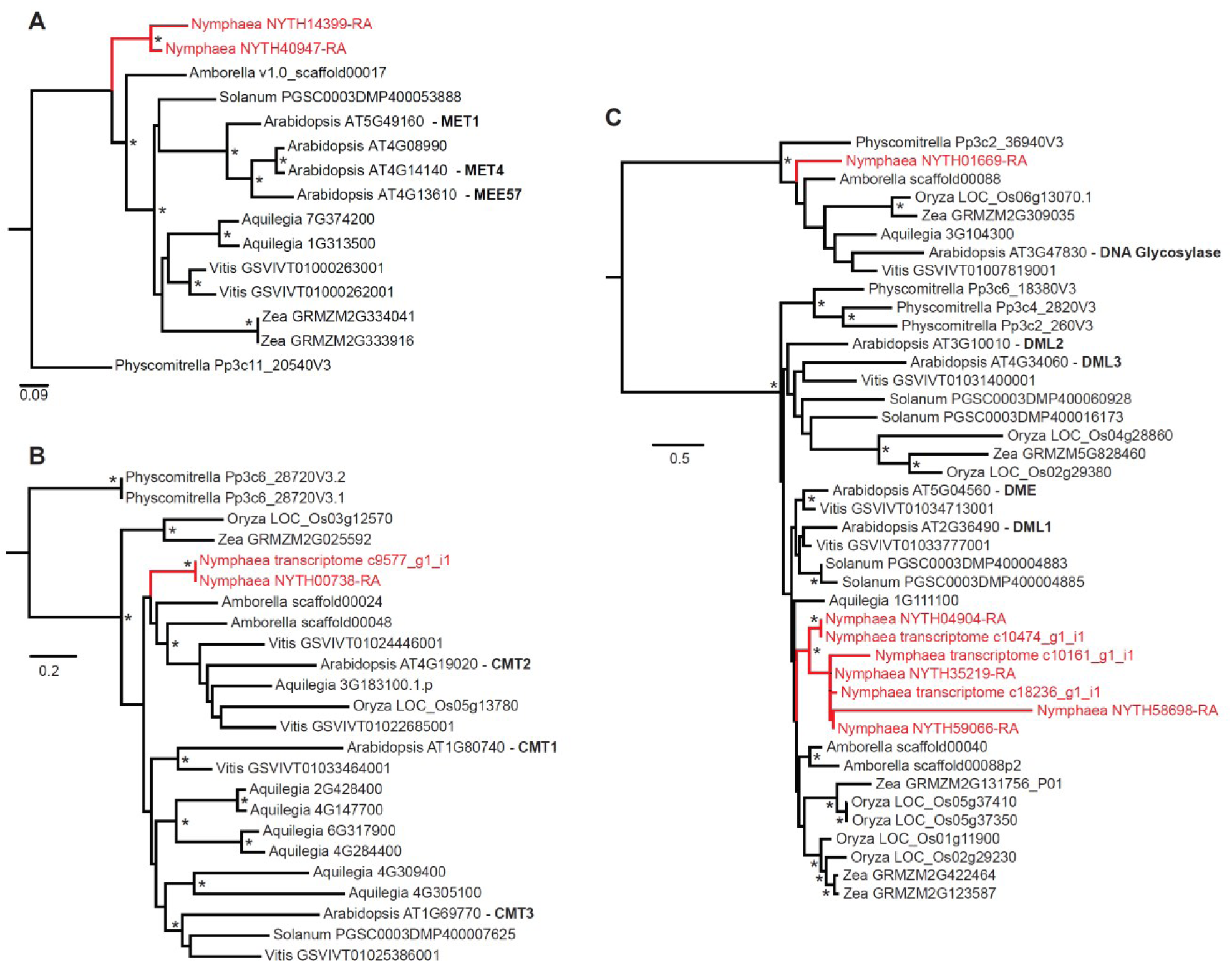
Gene family evolution for imprinting-related genes involved in the regulation of DNA methylation patterns. All gene family phylogenies were calculated with RAxML, from trimmed amino acid alignments. Bootstrap support (n=100) > 0.75 indicated by an asterisk. For each included sequence, the organism (genus name) and transcript identifier are noted. The gene names for *Arabidopsis* copies of interest are included in bold text. *Nymphaea* sequences (from transcriptome and genome assemblies) and sequence lineages are colored red. A) MET gene family. B) CMT gene family. C) DME (and DML) gene family.

**Figure 9:**
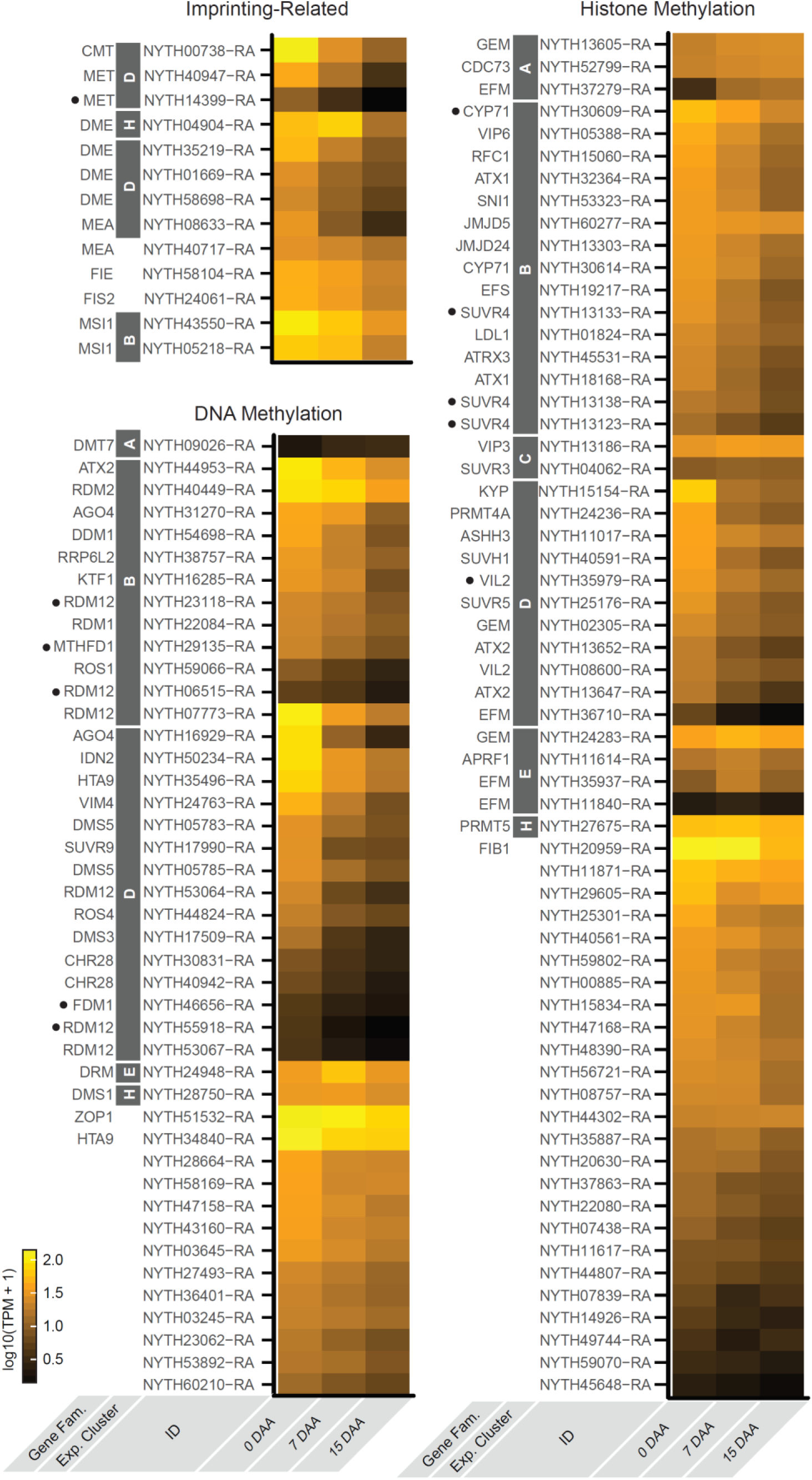
Expression of N. thermarum transcripts putatively involved in DNA and histone methylation. Transcripts putatively related to imprinting are noted separately from all other transcripts putatively involved in DNA or histone methylation. For each transcript, the following information is included (from left to right): gene family the transcript is associated with or the gene name of the most closely-related *Arabidopsis* homolog; whether the transcript is present in a DE expression pattern cluster; transcript identifier; expression at each of the sampled stages of reproductive development.

DME, on the other hand, removes certain types of methylation marks from DNA. We find that angiosperm DME genes were divided into two poorly-supported clades: one with DME and DML1, and one with DML2 and DML3 (Figure 7C). The *N. thermarum* DME homologs formed a single well-supported clade, suggesting clade-specific gene duplication events, that was placed (with poor support) within the [DME, DML1] clade. Three of the four *N. thermarum* DME homologs were in expression Cluster D (decrease after 0 DAA). The fourth and most highly expressed DME homolog, while in expression Cluster H (minimum at 15 DAA), did in fact display a significant increase in expression after 0 DAA (Figure 9).

*N. thermarum* homologs were also identified for all components of PRC2, and all were expressed during reproductive development. Angiosperm homologs of MEA formed two well-supported clades: one with MEA and SWN, and one with CLF (Figure 8A). Two *N. thermarum* MEA homologs were identified, with one present in each of the MEA clades. The *N. thermarum* homolog within the MEA/SWN clade was expressed during reproductive development, but not differentially expressed; the *N. thermarum* homolog of CLF associated with expression cluster D (decreased expression after 0DAA) (Figure 9). Two *N. thermarum* homologs of MSI1 were identified, and both associated with expression cluster B (consistent decreased expression) (Figures 8, 9). The FIE gene family appeared to be relatively simple, with little indication of gene duplications outside of monocots – one copy of FIE was identified from *N. thermarum* (Figure 8C). Angiosperm FIS2 genes formed two well-supported clades: one appeared to be specific to *Arabidopsis* (and included VRN2 and FIS2), while the other included EMF2. Only one homolog was identified from *N. thermarum*, and its placement within the EMF2 clade was poorly supported (Figure 8D). The FIE and FIS2 homologs in *N. thermarum* were expressed during reproductive development, but their expression did not significantly change during the sampled time points (Figure 9).

**Figure 8:**
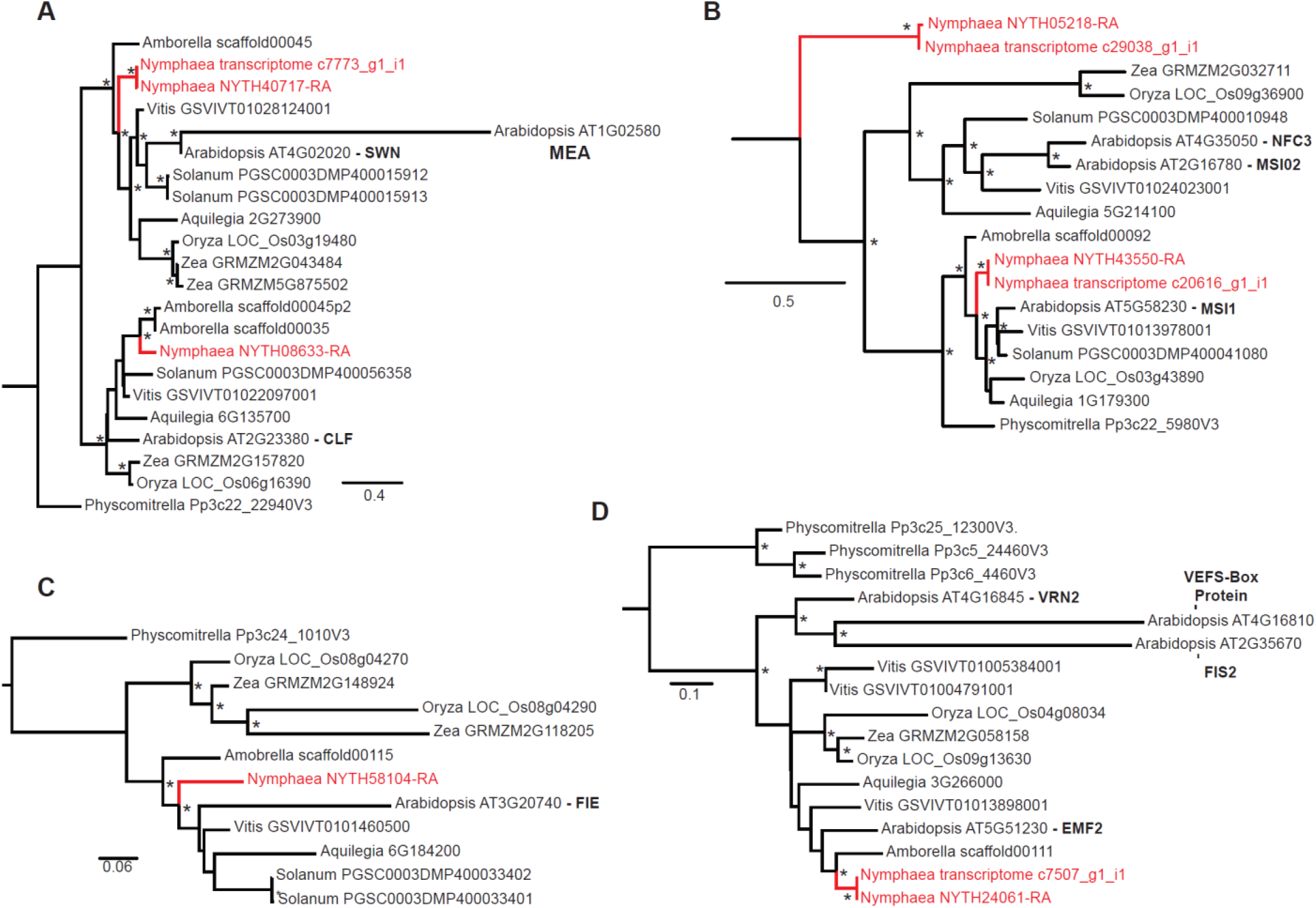
Gene family evolution for PRC2 components. All gene family phylogenies were calculated with RAxML, from trimmed amino acid alignments. Bootstrap support (n=100) > 0.75 indicated by an asterisk. For each included sequence, the organism (genus name) and transcript identifier are noted. The gene names for *Arabidopsis* copies of interest are included in bold text. *Nymphaea* sequences and sequence lineages are colored red. A) MEA (and CLF) gene family. B) MSI1 (and MSI02, NFC3) gene family. C) FIE gene family. D) FIS2 (and VRN2 and EMF2) gene family.

### Broader analysis of gene activity associated with DNA and histone methylation

We next used a broader approach to examine the expression of any gene that could be involved in regulation of DNA or histone methylation patterns (Figure 9). 121 loci in *Arabidopsis* thaliana are annotating as being involved in DNA or histone methylation; 125 putative homologs were identified from within the *N. thermarum* genome using pair-wise blast comparisons. 112 of the *Nymphaea* methyltransferase-related homologs were expressed in mature ovules or during seed development with a TPM >1, and of those 73 were significantly differentially expressed. Of the 112 putative DNA or histone methyltransferase-related homologs in *N. thermarum* expressed in mature ovules or developing seeds, 20 of them were not present in transcriptomes of root tips, leaves, young floral buds, or young ovules (Povilus 2020). 11 of the mature ovule/seed-development-specific homologs were differentially expressed, including putative homologs of RDM12, MTHFD1, MET, FDM1, CYP71, SUVR4, and ATX2; all were in the expression-pattern clusters that represented either general expression decrease (Cluster B), or decreased expression after 0DAA (Cluster D).

Most of the *Nymphaea* DNA and histone methylation-associated transcripts were in expression-pattern clusters that represented decreased expression at some point (Clusters B,D,H). Only 11 transcripts were among clusters that involved an increase in expression (A,C,E), including homologs of DTM7, DRM, GEM, CDC73, EFM, VIP3, SUVR3, and APRF1. Among the set of non-DE transcripts, a few were present with fairly high abundance, including homologs of ZOP1, HTA9, and FIB1.

## Discussion

### Overview

We leverage the ability of RNAseq datasets to move beyond a candidate gene approach, to broadly study seed development in the water lily *Nymphaea thermarum*, and specifically the processes involved in regulating imprinting-related and non-imprinting-related DNA and histone methylation. We find that all components of known imprinting mechanisms are expressed during reproductive development in *N. thermarum*, and that many other DNA or histone methylation regulators are differentially expressed. This indicates that not only is the epigenetic landscape likely to be dynamic during reproduction in *Nymphaea*, but that imprinting may also be occurring in this species. Comparisons with patterns of gene expression during reproductive development in other angiosperms suggests that the current model for how imprinting is regulated, perhaps best studied in *Arabidopsis*, is likely a mix of deeply conserved and eudicot-specific processes. Finally, we are able to suggest that the function of several histone-methylation genes merit further investigation during seed development in not only *N. thermarum*, but any model system.

### Patterns of gene expression during reproductive development

We find that a large proportion of genes is expressed during reproductive development in *N. thermarum*: 74% of the total number of genes predicted from the *N. thermarum* genome. Furthermore, 56% of the expressed transcripts are differentially expressed. The proportion of genes expressed, either differentially or not differential, during reproductive development is similar to what has been described in other species (Chen 2014). The number of unique DE transcripts in *N. thermarum* suggests that the transitions from female gametophyte maturation, through fertilization, and into mid-seed development require substantial transcriptional reprogramming (Figure 2B). While female gametophyte and ovule maturation involve relatively high numbers of unique genes (1,148), the expression of almost as many genes (916) appears to carry over into early seed development. 7 and 15 DAA shared the expression of far fewer genes (339; other than those shared by all three stages) – a surprising result given that 7 and 15 DAA are understood to share more developmental processes than 0 and 7 DAA. However, a previous study provided evidence for a lingering maternal influence on early seed development in *N. thermarum* (Povilus 2018), which is congruent with the relatively large number of transcripts shared between 0 and 7 DAA. Furthermore, when transcript expression pattern (not just presence/absence of transcripts) is taken into account, 7 and 15 DAA samples were more similar to each other than either were to 0 DAA samples (Figure 3A).

When DE transcripts are clustered by expression pattern, there are clear similarities among the enriched putative molecular functions for clusters that represent either increases or decreases in expression. As could be predicted by the onset of nutrient import and storage after fertilization, most of the “increased-expression” clusters were enriched for transporter activities, and a homolog of WAXY starch synthase 1 was among the 10 most highly expressed genes at 7 and 15 DAA. However, we also note patterns of gene expression associated with the onset of embryogenesis and/or endosperm development: highly-DE transcripts in Cluster E (maximum expression at 7 DAA) were enriched for transcription factor activity. Furthermore, the set of DE transcription factors in Cluster C and E (both involve expression increased between 0 and 7 DAA) were enriched for (respectively) WRKY and MYB genes, which have been associated with embryo and endosperm development in both *Arabidopsis* and *Zea mays* (Lagacé 2004)(Luo 2005)(Dubos 2010) (Wickramasuriya 2015).

Expression pattern clusters that represent a decrease in expression were enriched with an altogether different set of molecular functions. DNA/chromatin binding, transcription factor activity, and control of DNA polymerase are featured prominently in both the all-DE and highly-DE datasets. Intriguingly, expression Cluster D (decrease after 0 DAA) was enriched for GRF and ZF-HD transcription factors, which are associated with, among other things, cell division and floral development (Omidbakhshfard 2015). We attribute the pattern of decreased DNA-modification or transcription-regulation functions to either the cessation of cell proliferation and differentiation associated with ovule development, and/or the transition from floral development programs to seed development programs.

### Evidence for dynamic epigenetic landscape during reproductive development

In *N. thermarum*, 125 genes putatively share homology with *Arabidopsis* genes involved in DNA or histone methylation. A remarkable 89% of these *N. thermarum* homologs are expressed in mature ovules or during seed development (at TPM > 1), with 58% being differentially expressed, suggesting a dynamic epigenetic landscape during reproductive development in this species. Among the gene families known to specifically regulate imprinting-related methylation patterns, MET and CMT homologs were recovered in expression Cluster D (decreased expression after 0 DAA), as were three-quarters of the DME homologs. Furthermore, one of the MET homologs appears to be specifically expressed during seed development. The fourth *N. thermarum* DME, while associated with expression cluster H (decreased expression after 7 DAA), did in fact display a significant increase in expression after 0 DAA. All components of PRC2 were expressed during reproductive development.

Many components of the RdDM pathway were present during the sampled developmental stages in *N. thermarum*. Most fell into expression-pattern clusters B and D (consistent decrease in expression, or decreased expression after 0 DAA). Interestingly, most of the DNA or histone methylation-related homologs expressed only during seed development are components of the RdDM pathway (RDM12, FDM1), are known to be involved in chromatin remodeling (CYP71), and/or have been specifically tied to transposon repression (SUVR4, MTHFD1). In addition, DRM, an important component of the RdDM pathway, showed increased expression after fertilization. Homologs of several genes involved in histone methylation (GEM, CYP71, EFM, VIP3, SUVR3, APRF1) showed increased expression after fertilization. Several of these genes have not been previously linked to seed development in any species; we therefore suggest that their role during sexual reproduction deserves further investigation in *N. thermaurm* and other angiosperms, such as *Arabidopsis*, rice, and maize.

Altogether, our data suggests that DNA methylation patterns are being established, maintained, and removed before fertilization in *N. thermarum*. After fertilization, gene activity related to DNA methylation maintenance in the CG and CHG context (CMT, MET) decreases, while the expression of some genes involved in DNA demethylation (DME) and CHH-context de novo methylation (DRM and other RdDM components) increases. The components of PRC2, which establish loci-specific H3K27 methylation patterns associated with imprinting, all decrease in expression over time. By the time that the embryo typically initiates cotyledons at 15DAA, the expression of nearly all DNA and histone methylation-related genes has decreased in whole seeds, relative to their levels in pre-fertilization ovules.

### Comparison of imprinting-related DNA and histone methylation activity with other angiosperms

DNA and histone methylation during sexual reproduction has been studied for a small handful of distantly-related angiosperms, in particular the eudicot *Arabidopsis* and the monocot *Oryza* (rice)(Köhler 2012). Importantly, DNA and histone methylation have been shown to be dynamic during seed development in every taxon which has been studied. Although there is wide-spread evidence for parent-of-origin effects on seed development (Haig 1991), the molecular/genetic evidence for imprinting, which depends on patterning of DNA and histone methylation, is less consistent (Gleason 2012). It must be noted, however, that developmental stage sampling is inconsistent in many studies of gene expression during seed development (due in part to fundamental differences in how seeds develop), so comparisons should be approached with caution. In addition, complex, lineage-specific histories of gene duplication and loss can make it difficult to assess specific homology relationships within gene families.

A summary of expression patterns for genes related to imprinting in ovules and seeds of *Nymphaea*, monocots (mostly *Oryza*), and *Arabidopsis* is presented in Figure 10. CMT, and MET homologs all show decreased expression after fertilization in *Nymphaea*. While decreased expression of CMTs is similar to what is seen in *Arabidopsis* and rice, the expression pattern of the *Nymphaea* METs is the opposite of those in *Arabidopsis* and rice (Sharma 2009)(Julien 2012). The comparison for DME is more complex – one DME copy in *Nymphaea* shares the expression pattern with one barley DME homolog (Kapazoglou 2013). The expression of the second DME copy in *Nymphaea* is more similar to most of the rice DME homologs and to the *Arabidopsis* DME (Choi 2002)(Jiang 2016). All copies of PRC2 components decrease in expression after fertilization in *Nymphaea*, while expression patterns of individual components show more variation in *Arabidopsis* and rice (Baroux 2006)(Anderson 2013)(Nallamilli 2013).

**Figure 10:**
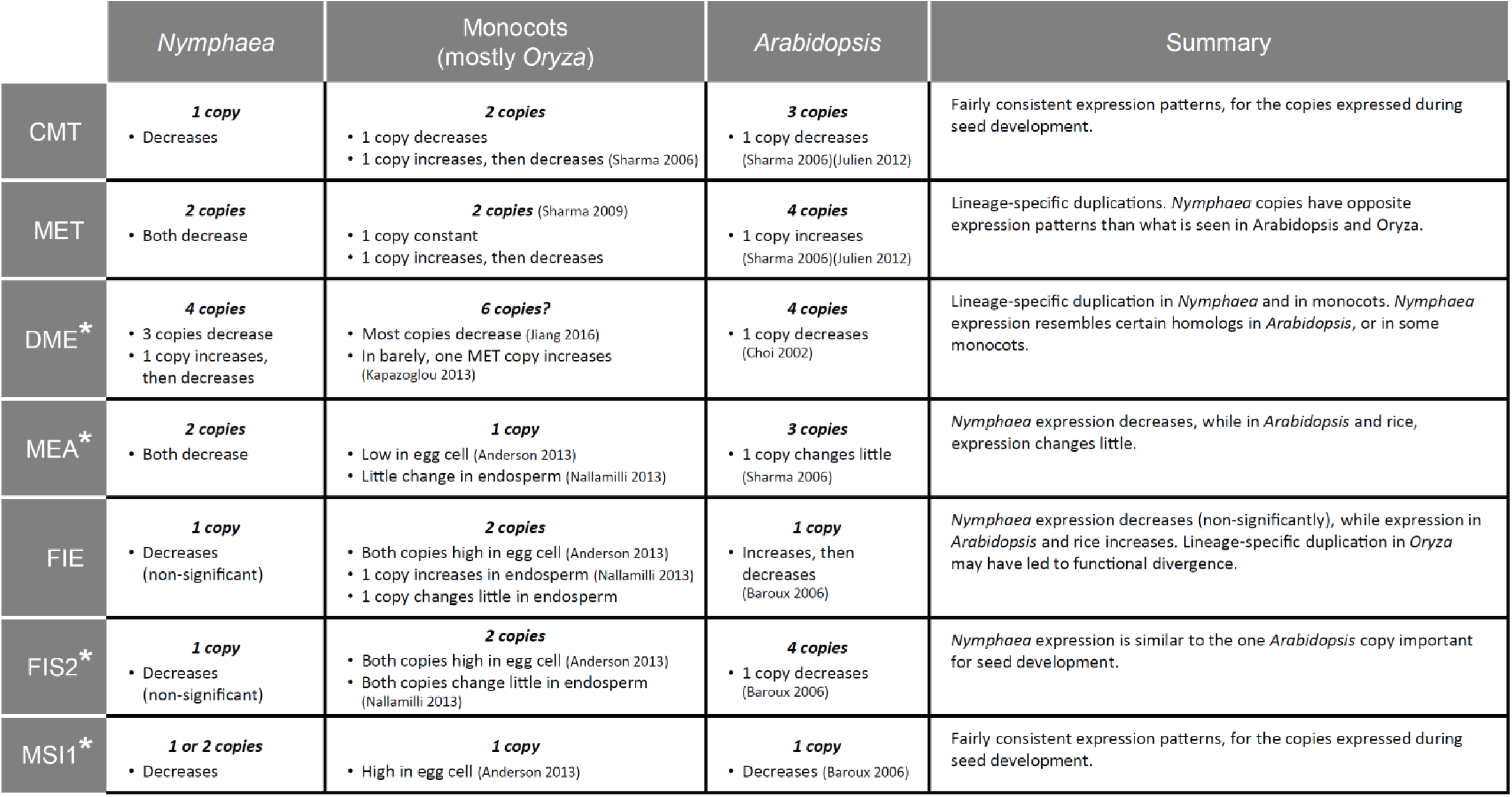
Summary and comparison of imprinting-related methylation regulators in *Nymphaea*, monocots (mostly *Oryza*), and *Arabidopsis*, and their expression before and after fertilization. For each gene family of interest, the number of copies in each species is reported, as well a brief summary of their relative expression before and after fertilization. An asterisk next to the gene family name indicates that a broad definition of gene family was used when assessing copy number (for example, the MEA* gene family includes both the MEA and CLF subfamilies).

Based on the complexity of DNA methylation-related activity, we conclude that DNA methylation is likely as dynamic during reproductive development in *N. thermarum*, as it is in other angiosperms. If imprinting occurs, however, its regulation is likely different than what is known for *Arabidopsis* – particularly with respect to the roles of MET and PRC2. Overall, maintenance of CG methylation by MET may be relatively less important after fertilization. In contrast, de novo CHH methylation by DRMs and other RdDM components may be relatively more important-whether or not they are related to imprinted gene expression. Interestingly, there is little evidence that PRC2 as a whole is a major regulator of seed development in rice (Luo 2009). Yet imprinting occurs in monocots, and impacts endosperm development – other molecular machinery must be responsible for regulating and responding to imprinting-related methylation patterns. Until the functions of the *Nymphaea* PRC2 homologs can be determined, we suggest that the role of the PRC2 as a whole in regulating imprinting may represent a derived condition within eudicots. Importantly, *N. thermarum* homologs of all of the genes known to be involved in imprinting via DNA or histone methylation are expressed in mature ovules or developing seeds. While parental-allele-specific RNA expression data is required for verification, our results indicate it is possible that imprinting via regulation of DNA and histone methylation may be occurring in this species.

## Declarations

### Availability of data and material

Raw sequence data and assembled transcriptomes of *N. thermarum* have been submitted to the National Center for Biotechnology Information (NCBI) database under BioProject PRJNA718528. Biological material and all other data are available as Supplemental Data, or from the corresponding authors upon request.

### Code availability

Not Applicable

### Authors’ contributions

R.A.P and W.E.F. conceived of original premise of the project. R.A.P. grew plant samples, performed experiments, and analyzed data. R.A.P. wrote the manuscript with input from W.E.F.

*Additional declarations for articles in life science journals that report the results of studies involving humans and/or animals*

Not applicable

### Ethics approval (include appropriate approvals or waivers)

Not applicable

### Consent to participate (include appropriate statements)

Not applicable

### Consent for publication (include appropriate statements)

All authors have given consent to publish this work.

## Acknowledgements

We acknowledge support from the National Science Foundation: IOS-0919986 awarded to W.E.F., and DEB-1500963 and IOS-1812116 awarded to R.A.P.. We thank the Botanische Gärten der Universität Bonn for providing original plant material for propagation.

