## Supplementary material for "Transcriptomes across fertilization and seed development in the water lily *Nymphaea thermarum* (Nymphaeales) reveal dynamic expression of DNA and histone methylation modifiers": Supplemenary Information (materials and methods and tables)

### Supplementary Materials and Methods:

RNA extractions:

For RNA extractions, 30-50 mg of tissue was ground in liquid nitrogen with a mortar and pestle before being added to 0.4 mL extraction buffer (100 mM Tris-HCL (pH 9), 2% (v/v) 2-Mercaptoethanol, 2% (w/v) PVP40). 40 ųl 10% SDS was added and samples were centrifuged at 12k g for 10 minutes at 4 C. The aqueous phase was transferred to a new tube with 2 volumes of TRIzol (Invitrogen). 1/5 volume chloroform was added, samples were centrifuged at 12k g for 10 min at 4 C, and the aqueous phase was transferred to a new tube. TRIzol extraction was repeated as necessary, up to two more times. An equal volume of cold isopropanol was added to the final aqueous phase, and samples were precipitated at -20 C for 60 min. Pellets were resuspended in $H_20$, and 2 volumes of saturated acid-phenol:chloroform (5:1) were added. Samples were centrifuged at 12k g for 20 min at 4 C, and the aqueous phase was transferred to a new tube. The acid-phenol extraction was repeated as necessary, up to two more times. If necessary, an additional hot (60 C) acid-phenol phase extraction was performed. For 15 DAA, samples were further purified with a column-based clean-up, using RNeasy Plant Mini Kit (Qiagen) spin columns, according to manufacturer protocol.

**Supplementary Table 1**: Number of reads and proportion of reads mapped for each sample, and for all samples.

| Sample | lane 6 | lane 7 | lane 8 | total # reads (paired-end) | reads processed (# read pairs) | reads pseudoaligned (# read pairs) | % reads aligned |
|---|---|---|---|---|---|---|---|
| 1 | 28,513,314 | 28,648,800 | 28,702,606 | 85,864,720 | 42,932,360 | 32,536,875 | 0.757863649 |
| 2 | 33,267,596 | 33,432,100 | 33,489,220 | 100,188,916 | 50,094,458 | 38,815,246 | 0.774841121 |
| 3 | 23,998,898 | 24,115,398 | 24,156,412 | 72,270,708 | 36,135,354 | 28,344,177 | 0.784389078 |
| 4 | 22,002,490 | 22,097,360 | 22,143,926 | 66,243,776 | 33,121,888 | 26,898,197 | 0.812097336 |
| 5 | 25,246,136 | 25,382,372 | 25,411,120 | 76,039,628 | 38,019,814 | 31,515,995 | 0.828936065 |
| 6 | 23,799,238 | 23,913,086 | 23,994,098 | 71,706,422 | 35,853,211 | 29,543,087 | 0.824001147 |
| 7 | 25,719,940 | 25,841,438 | 25,881,410 | 77,442,788 | 38,721,394 | 31,750,444 | 0.819971616 |
| 8 | 24,241,300 | 24,331,190 | 24,425,946 | 72,998,436 | 36,499,218 | 29,397,934 | 0.805440106 |
| 9 | 24,367,252 | 24,484,652 | 24,513,926 | 73,365,830 | 36,682,915 | 26,926,752 | 0.734040684 |
| 10 | 26,928,802 | 27,053,360 | 27,109,876 | 81,092,038 | 40,546,019 | 28,736,670 | 0.708742084 |
| 11 | 27,721,958 | 27,863,756 | 27,890,298 | 83,476,012 | 41,738,006 | 28,537,801 | 0.683736569 |
| 12 | 26,195,536 | 26,316,798 | 26,366,644 | 78,878,978 | 39,439,489 | 24,840,072 | 0.62982743 |
| | | | | Total number reads: | | Total number of reads pairs aligned (and total number reads aligned): | Total % reads aligned: |
| | | | | 939,568,252 | | 357,843,250 (715,686,500 total) | 0.761718479 |

**Supplemental Table 2:** Comparison of linear models, testing for the effect of time (daa) on the mean TPM values (per time point) of the 0.1% most highly expressed transcripts

Analysis of Variance Table

Model 1: log2(meanTPM + 1) ~ 1

Model 2: log2(meanTPM + 1) ~ daa

| | Res.Df | RSS | Df | Sum of Sq | F | Pr(>F) |
|---|---|---|---|---|---|---|
| Model 1 | 74 | 73.014 | | | | |
| Model 2 | 72 | 58.970 | 2 | 14.043 | 8.5732 | 0.0004573 |
